# Gene structure heterogeneity drives transcription noise within human chromosomes

**DOI:** 10.1101/2022.06.09.495447

**Authors:** Michael Chiang, Chris A. Brackley, Catherine Naughton, Ryu-Suke Nozawa, Cleis Battaglia, Davide Marenduzzo, Nick Gilbert

**Affiliations:** SUPA, School of Physics and Astronomy, University of Edinburgh, Peter Guthrie Tait Road, Edinburgh, EH9 3FD, UK; MRC Human Genetics Unit, Institute of Genetics and Cancer, University of Edinburgh, Western General Hospital, Crewe Road South, Edinburgh, EH4 2XU, UK

**Keywords:** Genome organisation, chromatin modelling, mechanistic models, polymer physics, transcription

## Abstract

Classical observations have long suggested there is a link between 3D gene structure and transcription^1–4^. However, due to the many factors regulating gene expression, and to the challenge of visualizing DNA and chromatin dynamics at the same time in living cells, this hypothesis has been difficult to quantitatively test experimentally. Here we take an orthogonal approach and use computer simulations, based on the known biophysical principles of genome organisation^5–7^, to simultaneously predict 3D structure and transcriptional output of human chromatin genome wide. We validate our model by quantitative comparison with Hi-C contact maps, FISH, GRO-seq and single-cell RNA-seq data, and provide the 3DGene resource to visualise the panoply of structures adopted by any active human gene in a population of cells. We find transcription strongly correlates with the formation of protein-mediated microphase separated clusters of promoters and enhancers, associated with clouds of chromatin loops, and show that gene noise is a consequence of structural heterogeneity. Our results also indicate that loop extrusion by cohesin does not affect average transcriptional patterns, but instead impacts transcriptional noise. These findings provide a functional role for intranuclear microphase separation, and an evolutionary mechanism for loop extrusion halted at CTCF sites, to modulate transcriptional noise.

---

Transcription – the copying of a DNA sequence into an RNA molecule – is a fundamental process, and the set of genes which are transcribed is essential to determine cell identity. Yet, the mechanisms underlying the selection of such active genes remains ill-understood. In human cells, DNA is tightly wrapped around histone proteins to form a composite polymeric material known as chromatin. Three-dimensional (3D) chromatin folding at the level of a gene and its surrounding regulatory domain (at the scale of 50-500 kilo-basepair, or kbp) is collectively termed “large-scale chromatin structure”. Long-standing experimental observations suggest a link between large-scale chromatin structure and the regulation of transcriptional output. For example, transcription often requires the formation of a 3D loop between the promoter of a gene and its enhancers^4^. However, experiments perturbing 3D structure or transcription genome-wide provide conflicting and unclear results^8–10^: for example, cohesin degradation leads to a dramatic perturbation of 3D structure genome-wide, but has little impact on gene expression^8^. This puzzling result shows that key insights into the mechanistic nature of the relationship between 3D structure and transcription are missing. Is 3D gene structure a consequence of other processes, or does it determine them? Can 3D chromatin structure provide the elusive mechanism through which cells determine which genes to activate and which to silence? This lack of clarity exists because experiments only probe the structure-transcription link indirectly, as it is impossible to follow chromatin dynamics and transcription simultaneously in the same cell.

## 3D gene structure genome-wide

To overcome these experimental limitations, we developed a coarse-grained polymer model to predict the 3D structure of chromatin in human GM12878 lymphoblastoid cells (see Methods for details). Coarse-grained models^11–14^ are aimed at building simplified representations of complex systems while retaining the main biophysical interactions. Here, we used an evolution of the “highly predictive heteromorphic polymer”, or HiP-HoP, framework^5^ which was further modified to consider both active and inactive chromatin, making it suitable for genome-wide studies. HiP-HoP is a mechanistic model which combines three main ingredients (Fig. 1a, and Extended Data Fig. 1). First, it includes multivalent chromatin-binding proteins, or factors, which can cluster through the bridging-induced attraction^6^, leading to intranuclear microphase separation. Second, it incorporates loop extrusion by cohesin or other structural maintenance of chromosome (SMC) proteins^7^, which is necessary to explain the formation of convergent CTCF loops in mammalian genomes^15^. Third, it accounts for chromatin heteromorphism, which is required to recapitulate the decompaction of highly active genes^5,16^.

**Fig. 1:**
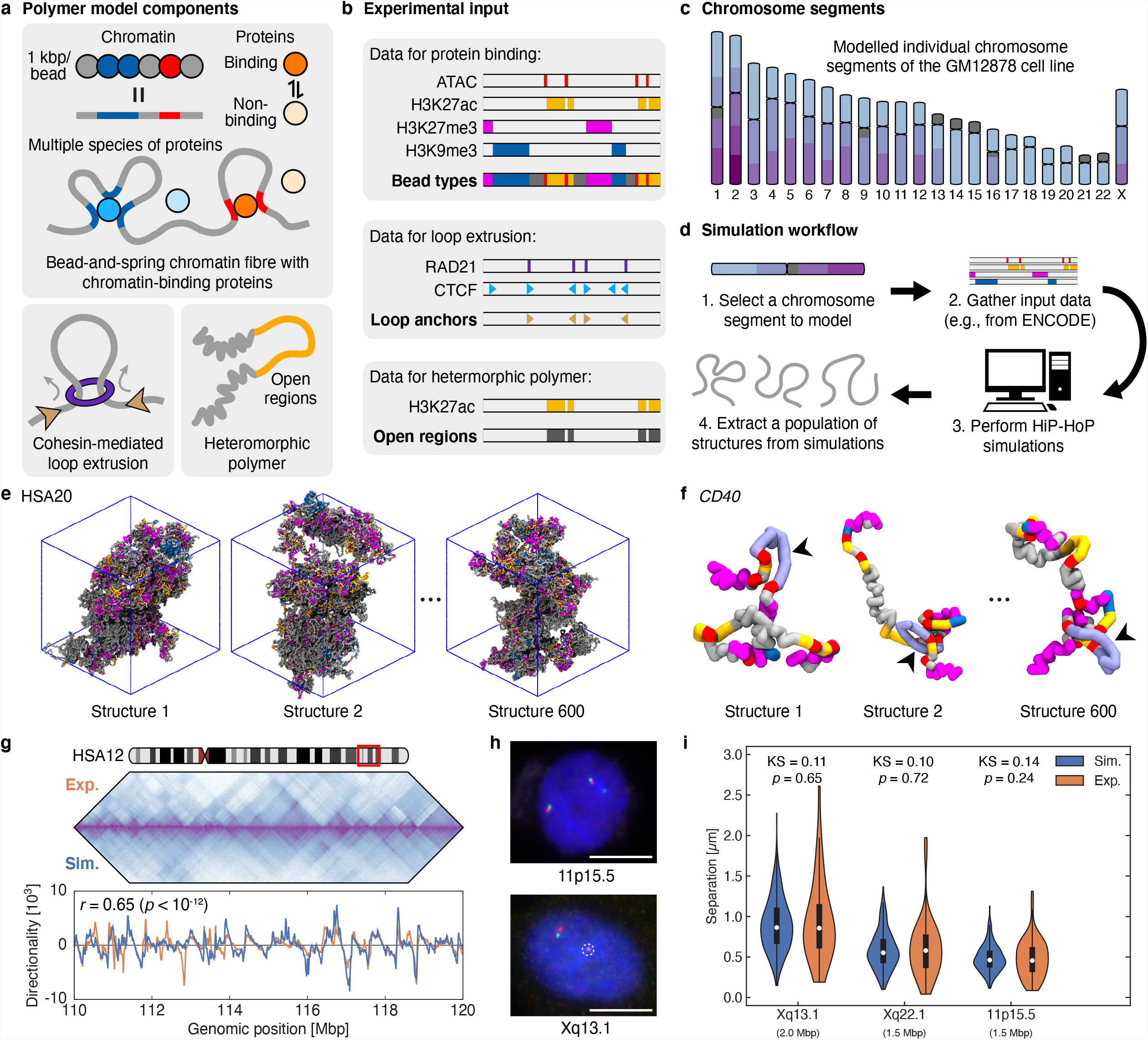
Genome-wide simulations of 3D chromatin structures using the HiP-HoP framework. **a**, Key ingredients of the framework. The chromatin fibre was modelled as a bead-and-spring polymer chain, with each bead representing 1 kbp of chromatin. Chromatin binding proteins, modelled as multivalent, spherical beads that can switch between a binding and non-binding state, were incorporated to bridge between specific chromatin loci. Loop extrusion was assumed to be mediated by SMC complexes, such as cohesin^7^, and to halt at convergent CTCF binding sites. Local variation in chromatin compaction was incorporated into the model by including extra folding to regions depleted of the H3K27ac mark (see Methods). **b**, Experimental epigenetic inputs necessary for HiP-HoP simulations. ATAC-seq peaks and ChIP-seq peaks for H3K27ac, H3K27me3, and H3K9me3 were used to classify the type, or flavour, of individual chromatin beads, such that they were bound by different proteins. ChIP-seq peaks for CTCF and Rad21, a subunit of the cohesin complex, were used to determine the positions of barriers (or anchors) against loop extrusion. H3K27ac ChIP-seq marks open or disrupted chromatin fibres. **c**, Simulations were performed for the entire genome of the GM12878 human lymphoblastoid cell line, with chromosomes divided into segments and simulated individually, as indicated by the partitions within the ideograms. **d**, Illustrations showing the typical workflow for running HiP-HoP simulations. 600 independent structures were obtained for each chromosome segment. **e**, As an example, snapshots of the structures of HSA20, simulated as a single segment. **f**, Enlarged views of an example gene locus, *CD40*, located within HSA20. The model can resolve the 3D structure of individual gene loci at 1 kbp resolution across the entire human genome. **g**, An example comparison between contact maps from Hi-C experiments^15^ and those predicted using HiP-HoP (top), and between the directionality profiles extracted from these contact maps (bottom; see Methods and Extended Data Fig. 5a-d). Pearson’s *r* is shown for correlating the profiles. **h**, Representative images of two-colour FISH probes used to measure distances between selected chromosome segments. Scale bar is 10 μm, and white dashed circle marks the inactive X chromosome. **i**, Violin plots comparing the experimental^41^ and simulated FISH distance distributions for probes in **h** and those located on Xq22.1 (see Extended Data Fig. 5e for more examples). A two-sample Kolmogorov-Smirnov (KS) test was used to determine if the distributions were statistically different.

Chromatin was modelled as 1 kbp building blocks (or “beads”), which were “painted” depending on different chromatin types^17^, according to input epigenetic 1D data sets (Fig. 1a and Extended Data Fig. 1, see Methods for more details). Three types of multivalent chromatin-binding factors were considered: a generic active factor (modelling complexes of RNA polymerases and transcription factors), and two types of repressive/silencing factors, modelling polycomb-like and heterochromatin-binding (HP1-like) respectively. The positions of binding sites for these factors along the chromatin (Fig. 1b) were inferred from ATAC-seq data (for active proteins), H3K27me3 (for inactive polycomb-like proteins) and H3K9me3 (for HP1-like inactive proteins). The experimental input for loop extrusion used Rad21 and CTCF ChIP-seq data to identify binding sites for CTCF that act like loop anchors and halt cohesin-mediated extrusion in a binding motif orientation specific way (Fig. 1b), while regions marked by H3K27ac were considered to form more disrupted chromatin fibres (see Ref. ^5^). The genome was broken into 52 segments at gene deserts or chromosome arm boundaries (Fig. 1c and Extended Data Fig. 2), and polymer simulations informed by the epigenetic input data (Fig. 1b) were then run (Fig. 1d; see Methods for more details on the simulation protocol and force fields used). Each individual simulation iterated through 3×10^7^ timesteps, corresponding to approximately 25 min in real time, and the structure of each genome fragment was simulated independently 600 times, providing an ensemble of structures (Fig. 1e and Extended Data Fig. 3) from which data for individual transcription units were abstracted such as *CD40* (Fig. 1f), *BCL2* and *SERPINB8* from HSA18 (Extended Data Fig. 3c) or *DNMT1* and *LDLR* from HSA19 (Extended Data Fig. 3d). Data were either combined to measure average structural properties or contact statistics, analogous to averaging many cells in a population, or considered individually, akin to single-cell techniques. The structure data generated for each gene is stored as CIF files within the 3DGene database (https://3dgene.igc.ed.ac.uk; Extended Data Fig. 4), accessible through a web-tool enabling researchers to view or download the predicted 3D structures, contact matrix and transcriptional activity (see below) interactively for any human gene, or genomic region.

As the HiP-HoP framework does not require 3D structural data as an input, it was previously validated by comparing its output to existing Hi-C, Capture-C and FISH data^5,18^. To further validate 3DGene at a genome-wide scale, bead-bead interaction data at 1kbp resolution were averaged between simulations and used to generate contact maps which were compared quantitatively to Hi-C data from GM12878 cells^15^ using domain directionality (D) (Fig. 1g, Extended Data Fig. 5a and methods). This comparison led to a good agreement (Pearson’s *r* ∼ 0.65, *p* < 10^−12^, for the region shown in Fig. 1g and correlation ranged between 0.4 and 0.7, Extended Data Fig. 5b-d), which is notable due to the absence of fitting in the analysis. Furthermore, distances between beads were used to generate probability distributions which were compared to those from two-colour FISH data (Fig. 1h). FISH distributions were well matched, especially within a gene locus (Fig. 1i and Extended Data Fig. 5e; differences between experimental and simulated distributions were typically not statistically significant).

## Human genes display a large variation in structure and structural diversity

To characterise gene structure genome-wide, we interrogated the 3DGene dataset. First, gene promoters were identified as chromatin beads associated with both an ATAC-seq peak in the experimental input and a promoter annotation^19^ (Fig. 2a). Then, 3D interactions within a gene were analysed by examining contacts between the promoter and every other bead associated with an ATAC peak (or “ATAC bead”). ATAC beads with which the promoter was spatially proximate in at least 10% of the simulated structures were termed “ATAC partners”, and those which were proximate in >50% of the structures were designated “influential nodes” (Fig. 2a). Each promoter is uniquely associated with a gene locus, defined as the genomic region encompassing all its partners. The median size of a gene locus is 209 kbp (Fig. 2b), significantly smaller than the average ∼1 Mbp size of topologically associating domains (TADs)^20,21^. Human gene loci have a highly variable number of partners and influential nodes (exponentially distributed; see Fig. 2b). In selecting a sample of genes with 14 partners the two extremes correspond to *ZBTB5*, a housekeeping gene coding for a zinc-finger protein without any influential nodes, and *PTPN22*, a lymphoid-specific gene coding for an intracellular phosphatase with seven influential nodes (Fig. 2c,d). Intriguingly, gene ontology (GO) analysis suggests that genes with more influential nodes tend to be tissue-specific, while those with few or none are more likely to be tissue-invariant, or housekeeping: for instance, the subset of genes with more than one influential node is enriched in GO terms related to immune response (Extended Data. Fig. 6), indicating that the number of influential nodes within a locus is a structural feature with immediate relevance to transcription.

**Fig. 2:**
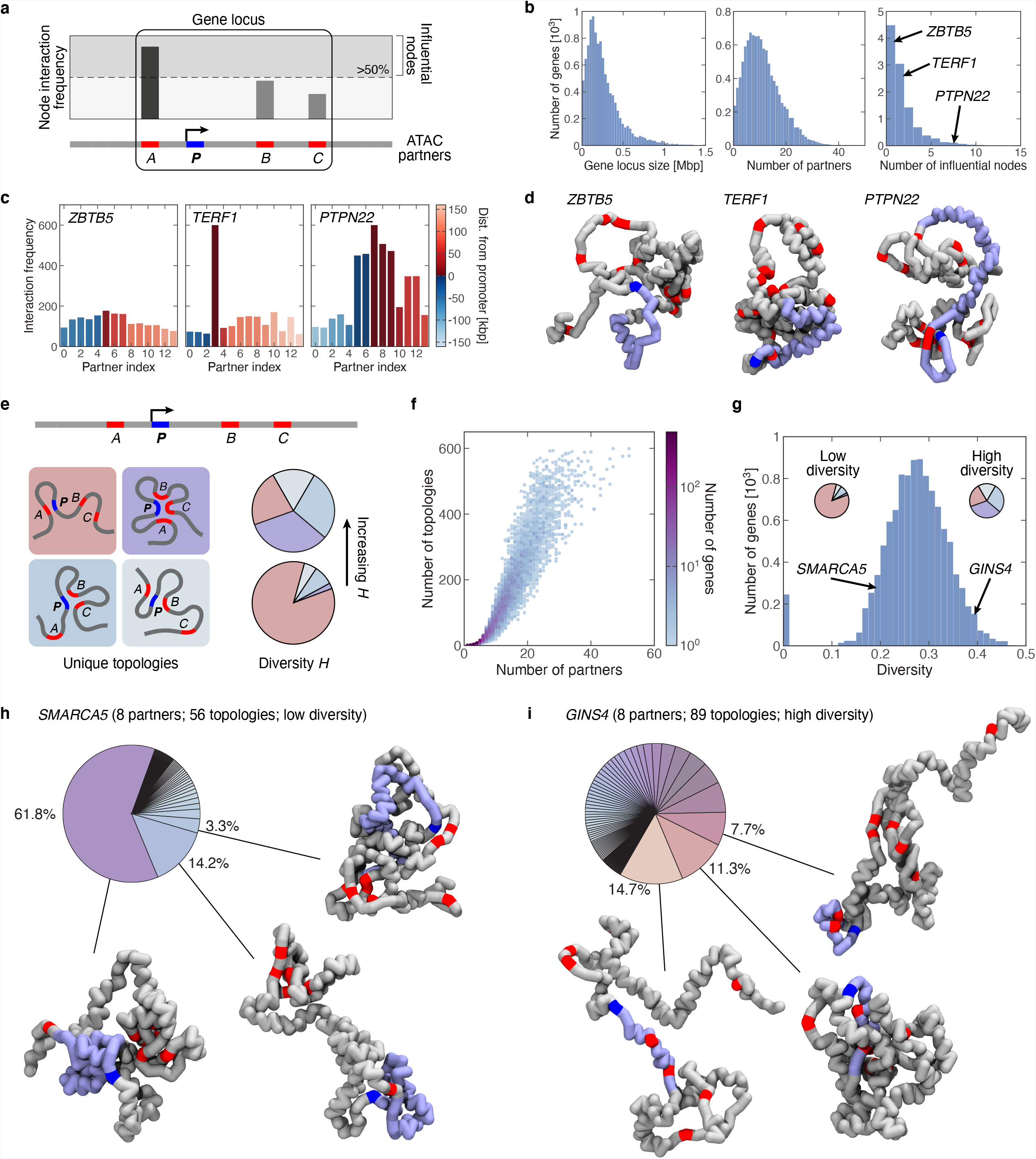
Large variability in the 3D structures and structural diversities of different gene loci. **a**, Schematics illustrating how ATAC partners and influential nodes are determined for each promoter. In our notation, the chromatin stretch including all partners of a promoter defines its gene locus. **b**, Histograms showing the distributions of size, number of partners and number of influential nodes for loci genome-wide. **c**, Interaction frequencies showing the number of structures where each ATAC partner in the locus contacts the promoter for three example loci with 14 partners and zero (*ZBTB5*), one (*TERF1*), and seven (*PTPN22*) influential nodes. The colour scale indicates the distance between each partner and the promoter. **d**, Examples of 3D structures for the *ZBTB5, TERF1*, and *PTPN22* loci. **e**, An illustration explaining the identification of topologies in a gene locus and its associated diversity *H* score. Here, with a population of five topologies, a higher *H* is achieved when the sampled conformations are distributed more evenly among them (i.e., more equal slices in the pie chart), whereas a lower *H* occurs when many conformations are assigned to one of the topologies (more unequal slices). **f**, A scatterplot showing the number of topologies against the number of partners for loci genome-wide. **g**, Histograms showing the distribution of structural diversities for loci genome-wide. **h**, Topology pie-chart and example structures for the most frequent three topologies for *SMARCA5*, a low-diversity locus with eight partners and 56 topologies. **i**, similar to **h** but for *GINS4*, a high-diversity locus with eight partners and 89 topologies.

To better classify gene structure, we defined topology as a unique subset of partners of a promoter which interacts with it for a given structure, and catalogued all observed interaction topologies of any gene locus. For example, the locus shown in Fig. 2e has one topology where the promoter *P* interacts with the ATAC bead *A* (top left) and another where *P* interacts with both *A* and *B* (bottom right). To quantify the structural diversity of a given locus, we used a concept borrowed from information theory, and defined diversity as the normalised Shannon entropy (*H*), measuring the relative share in the frequencies of detecting individual topologies (Fig. 2e and Methods). The number of observed topologies is highly correlated to the number of common partners for all gene loci, but shows striking variations between different gene loci with similar numbers of partners (Fig. 2f).

Across the genome there is large variability in diversity, indicative of structural heterogeneity at different gene loci (Fig. 2g). The *SMARCA5* locus, encoding a component of the chromatin remodelling and splicing factor RSF facilitating RNA Pol II transcription, has eight ATAC partners, and 56 different topologies (Fig. 2h). The most common one accounts for over 60% of all simulated structures, and concomitantly *SMARCA5* has low structural diversity (Fig 2g). In contrast, *GINS4*, encoding SLD5, a key component of the DNA replication complex, has the same number of partners, but 89 topologies, with the most common one accounting for just 14.7% of the simulated structures (Fig. 2i), corresponding to high diversity.

## Transcription factor phase separation drives transcriptional activity

The previous descriptive analysis suggested a correlation between influential nodes and function, and between diversity and transcription (Fig. 2). To understand the mechanisms leading to these correlations, we reasoned that transcription initiation requires binding of complexes of transcription factors (TFs) and polymerases at gene promoters (for genic transcription), or other ATAC beads (for non-genic transcription, such as at enhancers^22^). We therefore hypothesised that the transcriptional activity can be estimated by measuring the fraction of time within a simulation that there is an active protein (e.g., a TF) spatially proximate to a promoter (Fig. 3a). Within our framework, we can gather information from single cells (corresponding to single simulations), and aggregate results to obtain a distribution of transcriptional activities (Fig. 3a).

**Fig. 3:**
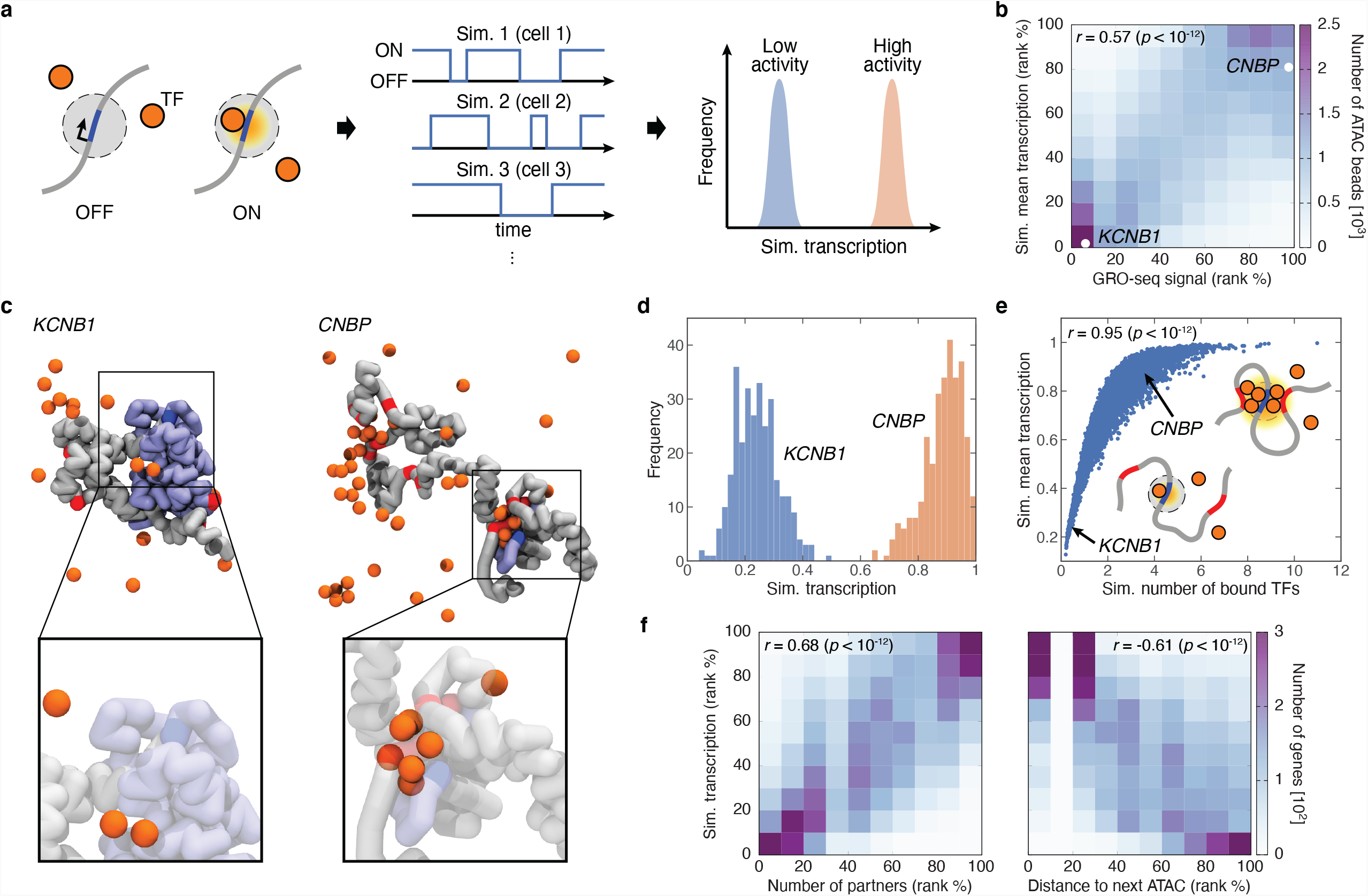
HiP-HoP simulations predict transcriptional activity of individual gene regulatory elements. **a**, Schematics explaining how TF binding at an ATAC bead in simulations is used to predict the transcriptional output of chromatin within the bead. Here, an ATAC bead (shown in blue) is envisaged to be in a poised state (OFF) unless a TF comes within its contact radius (3.5*σ*; grey circle), upon which it becomes activated (ON). From recording a time series of TF binding activity, the fraction of time where at least one TF associates with an ATAC bead is measured for each simulation run, and the average of these determines the predicted transcriptional activity. **b**, A heatmap showing the number of ATAC beads in bins according to their percentile rank in the TF binding probability (predicted transcriptional activity) against their rank in the GRO-seq signal. **c**, Examples of 3D structure for a lowly transcribed (*KCNB1*) and a highly transcribed (*CNBP*) locus. In *KCNB1*, an enlarged view of the structure near the promoter (in blue) shows only two TFs bound to the transcribed (light purple) region, whereas in *CNBP* there is a microphase separated cluster of chromatin-binding proteins near the promoter. **d**, Histograms showing predicted transcriptional activities for the promoters of the *KCNB1* and *CNBP* loci, calculated by aggregating all simulations (see Methods). **e**, A scatterplot of the mean predicted transcriptional activity versus the mean number of TFs bound to the promoter: the latter quantity measures the extent of microphase separation within the locus, which strongly correlates with transcription. **f**, Heatmaps showing the number of genes (promoters) in bins according to their percentile rank in predicted transcriptional activity against their rank in the number of partners (left) and the distance between promoter and the nearest ATAC bead (right). Spearman’s *r* is reported for all heatmaps and scatterplots.

To test whether transcription can be faithfully predicted from 3DGene data, TF binding probability of a given ATAC bead was correlated to transcriptional activity measured experimentally using GRO-seq, yielding a Spearman’s *r* ∼ 0.57 (*p* < 10^−12^; Fig. 3b). If transcriptional activity were purely determined biochemically, for instance by specific affinity between TFs and the promoter DNA sequence or epigenetic modifications, no significant correlations between expression data and 3DGene predictions would be expected, as these simulations do not specify sequences inside an ATAC bead. Instead, this correlation shows that the structural features of a gene determine transcription.

To further dissect the link between structure and transcription within the 3DGene simulations, we correlated predicted transcriptional activity (i.e. TF binding probability) and other structural variables. Previous studies suggest two main and extreme models for the structure of active genes: the decompaction and microphase separation model^23^. The first is based on FISH experiments which have shown that many eukaryotic genes decondense when activated^5,16^. The microphase separation (or transcription hub or factory) model instead envisages that active genes form clusters, or hubs, where promoters and enhancers come together, surrounded by loops of the intervening chromatin^24–28^. 3DGene structures show that both models apply, with active genes exhibiting either a decompacted structure due to enhanced acetylation, or a looped and phase separated one due to augmented enhancer-promoter interactions. Quantitatively, the predicted transcriptional activity correlates both with locus radius of gyration (Spearman’s *r* ∼ 0.66, *p* < 10^−12^), as would be expected from the decompaction model, and with number of TFs close to the promoter (*r* ∼ 0.95, *p* < 10^−12^; Fig. 3f), as predicted by the microphase separated one, which therefore appears more pervasive. As typical examples of this more prevalent model, two genes were examined in detail. The *KCNB1* locus is unlooped and has a low average transcriptional activity (Fig. 3c), whereas the ATAC beads in the *CNBP* locus cluster and form a microphase separated droplet, with a concomitant increase in transcription (Fig. 3d). Overall, the average predicted transcriptional activity increases sharply with the number of bound TFs, which quantifies the local extent of microphase separation (Fig. 3e). The emerging picture from the analysis of 3DGene structures is that protein-mediated bridging between ATAC beads close in sequence space (Fig. 3f) drives formation of microphase separated protein clusters, which are highly active transcriptionally.

## Structural diversity and loop extrusion determine transcriptional noise

By analysing the distribution of transcription activity over a set of independent simulations (analogous to different cells in a population) for each gene locus, transcription variability or “noise” can be measured (as the standard deviation of the distribution; Fig. 4a). Such noise, scaled by activity to give a coefficient of variation, significantly correlates with the analogous quantity computed from single-cell RNA-seq data (Fig. 4b; see Methods). To establish whether noise was related to 3D structures, all measured structural features of simulated gene loci were correlated with the transcriptional noise at their promoters. The first salient correlation emerging is that noise positively correlates with structural diversity (Fig. 4c). For instance, *GINS4* and *SMARCA5* (Fig. 2h-i), with high and low diversity, are respectively a high-noise and a low-noise locus (Fig. 4d). Therefore, a functional role of structural diversity is to give rise to noisier transcription, which could be advantageous for specific genes, such as developmental ones.

**Fig. 4:**
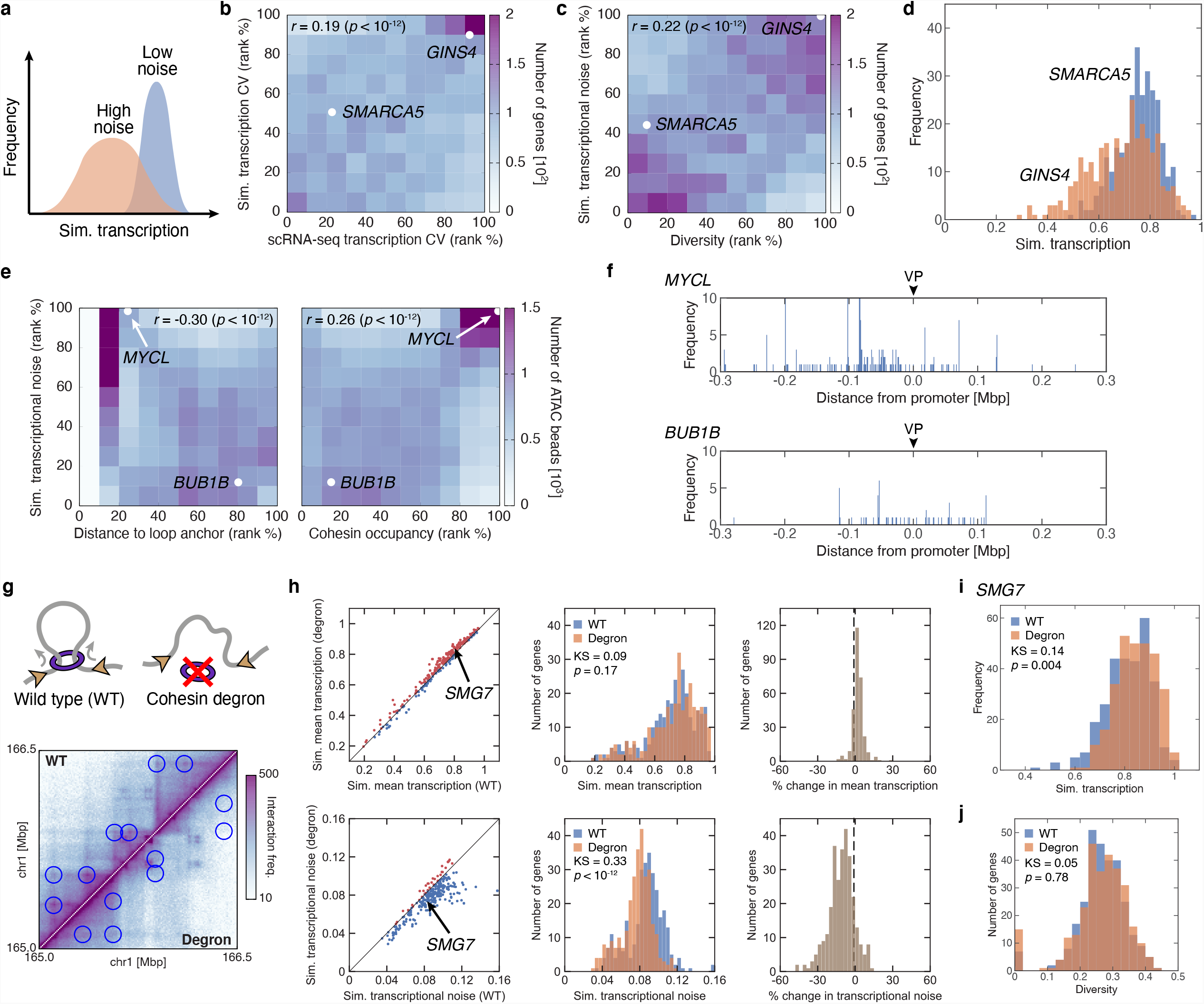
Structural diversity and loop extrusion determine transcriptional noise. **a**, Definition of transcriptional noise. For a given promoter or ATAC bead, we defined the transcriptional noise as the width (standard deviation) of the histogram of the simulated transcriptional activities over a population of cells (or many simulations). **b**, A heatmap showing the number of genes in bins according to their percentile rank in the coefficient of variation (CV) of transcription predicted from simulation against that from single-cell RNA-seq data (Ref. ^42^; see Methods). **c**, Similar to **b**, but for the rank in simulated transcriptional noise against that in structural diversity. **d**, Examples of histograms of simulated transcription for *GINS4* and *SMARCA5*, showing the former is noisier. **e**, Heatmaps showing the number of ATAC beads in bins according to their percentile rank in simulated transcriptional noise against their rank in the distance to the nearest CTCF bead (left) and the frequency of cohesin occupancy at the bead corresponding to the gene promoter (right). **f**, Examples of 4C-like profile for cohesin-mediated loops involving the promoter (i.e., the viewpoint, VP) for two loci, *MYCL* and *BUB1B*: the former is noisier and closer to a CTCF site. **g**, An illustration (top) explaining the setup for cohesin degron simulations: these were performed with all extrusion springs removed from the chromatin fibre. Degron simulations were performed for the chromosome segment HSA1:142.5-189.0 Mbp. A comparison of the contact maps generated from the wild-type (WT; upper triangle) and degron (lower triangle) simulations is shown in the bottom. Interaction frequencies are shown in log scale to aid visualisation. Circles highlight the disappearance of “dots” (which typically correspond to loop anchors) at TAD boundaries upon cohesin removal. **h**, Scatterplot and histograms comparing the simulated transcriptional activity and noise of each gene locus in the WT and degron conditions. Transcriptional activity is essentially unaffected, while noise decreases significantly in the degron [two-sample Kolmogorov-Smirnov (KS) test]. **i**, Histograms of the simulated transcriptional activity for *SMG7* showing a decrease in noise (smaller width in the distribution) in the degron condition compared to the WT. **j**, Histograms of structural diversities of gene loci in the WT and degron conditions, showing little detectable change (two-sample KS test). Spearman’s *r* is reported for all heatmaps.

A second finding is an anticorrelation between transcriptional noise associated with an ATAC bead, and genomic distance to the nearest CTCF or cohesin binding site (Fig. 4e, left). From this observation, we reasoned that there might be a relation between loop extrusion, which is halted at convergent CTCF sites, and transcriptional noise. Consistently, there was a positive correlation between noise and cohesin occupancy, which measures the density of loop-extruding factors (Fig. 4e, right). The relation between CTCF and transcriptional noise is also apparent by a more detailed analysis of the topologies and transcriptional output of example loci. For instance, *MYCL* is a noisy gene, where the promoter is close to a CTCF anchor. The pattern of cohesin-mediated loops to the promoter (Fig. 4f, top) shows that these are abundant and diverse. In contrast, *BUB1B*, a gene with low noise and a distal CTCF anchor, has only few cohesin-mediated loops involving the promoter (Fig. 4f, bottom). The relative scarcity of loop extrusion activity of the promoter, we speculate, leads to a smaller range of possible 3D structures, and, in turn, to less transcriptional noise.

This analysis strongly points to a relation between loop extrusion and transcriptional noise. Loss of cohesin, a putative loop extrusion factor^7^, has been studied experimentally using cohesin degrons^8^, and results in the loss of loop domains and visible structural changes; however, little change in the expression level of a gene in a bulk population has been observed. To test the idea of cohesin having a role in transcriptional noise, we considered how noise might change after cohesin degradation by performing additional simulations on a selected chromosome segment without loop extrusion (Fig. 4g). Consistent with experimental data, simulations showed a loss of loops between CTCF sites, observed as ‘dots’ in contact maps (Fig. 4g), but no substantial change in the average transcriptional activity of promoters (or more generally ATAC beads; Figs. 4h). Strikingly, instead, transcriptional noise, determined from simulations, markedly decreased after simulating a loss in cohesin (Fig. 4h-i), indicating that loop extrusion regulates transcriptional noise, rather than the average activity. Interestingly, the correlation between noise and structural diversity is not affected by this loss in cohesin, suggesting there are two separate pathways to control gene noise (Figs. 4j and 5).

**Fig. 5:**
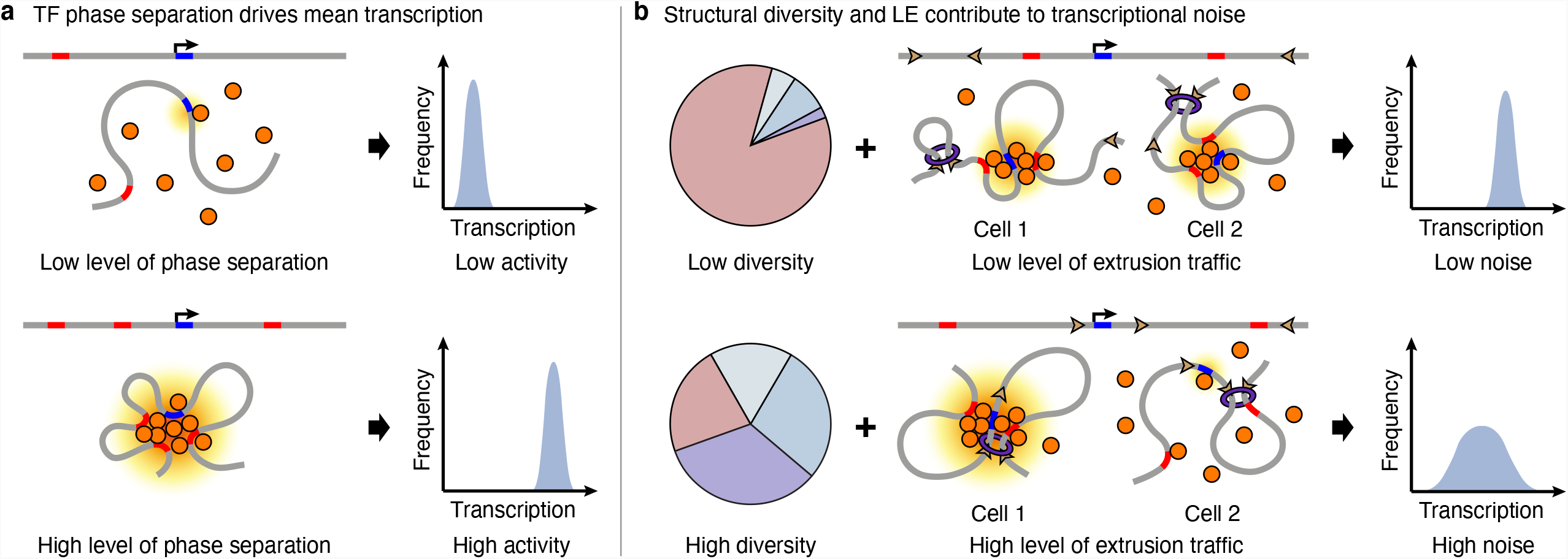
A mechanistic model for the link between 3D structure, transcription and gene noise. **a**, A mechanistic model for the relation between 3D structure and mean transcription. In a typical low-activity gene, the promoter is far from other regulatory elements (ATAC beads) and unable to form stable contacts with them. Instead, in a typical high-activity gene the high local density of regulatory elements promotes the formation of microphase separated droplets of active factors, driving transcription. **b**, Mechanisms contributing to transcriptional noise. Structural diversity associated with the patterning of ATAC-seq peaks and loop extrusion provide two complementary pathways to increase the heterogeneity of the 3D structure of a gene locus, determining its transcriptional noise.

## Discussion

Experiments involving auxin-inducible degrons of cohesin^8^ and RNA polymerase^10^ have pointed to a nuanced link between 3D gene structure and transcription. The link is difficult to dissect experimentally due to the challenges of collecting structure and transcription data concurrently, motivating our current alternative approach based on genome-wide mechanistic simulations of gene structure and function. We found that microphase separated droplets containing regulatory elements, arising through the bridging-induced attraction^6^, are a fundamental structural unit with a direct role in transcription, as larger droplets tend to be associated with higher transcriptional activity (Figs. 3e-f and 5a). Such structures are reminiscent of active transcriptional hubs previously proposed on the basis of experimental microscopy data^25^. These data sit well with the recent observation from enhancer mobilisation experiments that enhancer action depends on genomic distance, and that contacts determine transcriptional activity nonlinearly^29^. From our simulations, validated by comparison with experimental data (Hi-C, GRO-seq, FISH and single-cell RNA-seq), we also showed that there are two separate pathways through which structure determines gene noise. First, noise is controlled by structural diversity (Fig. 4b and 5b), which is in turn determined by the pattern of accessible elements (ATAC-seq peaks) in a specific gene locus. Another mechanism to regulate transcription noise is through loop extrusion driven by SMC proteins, which modulate structural heterogeneity (Fig. 4g-j and 5b). These results provide a framework to interpret results on cohesin degrons, which typically only find subtle effects on transcription^8^. As cohesin depletion nevertheless leads to a small decrease in the typical number of partners in a locus (median of 11 partners in the wild-type condition compared to 9 in degron), our results are also consistent with recent findings that cohesin is required for long-range enhancer-promoter interactions^30^.

As evolutionarily young genes are known to be noisier than older ones^31^, we speculate that loop extrusion, in conjunction with CTCF positioning, may have been exploited by evolution to regulate gene noise, which may be advantageous for certain developmental genes^32^, where higher noise implies higher tunability, but also generates cellular heterogeneity. What mechanism enables loop extrusion by SMC proteins to increase gene noise? We hypothesise that this is because cohesin-CTCF loops form stochastically, and are stable for minutes^33^, so that a different network of cohesin-CTCF loops form in a different cell (or simulation), and this increases the variability in chromatin topologies and contact probability for a given ATAC bead, thereby increasing noise (Figs. 4e, bottom and 5b).

Besides the link to transcriptional hubs and cohesin degron experiments, our results provide insight into other biological observations that are difficult to reconcile with a single existing model. First, ImmunoGAM suggests that chromatin topologies are cell-type specific^34^, at variance with TADs, which are largely cell-type insensitive^35^. Our results also suggest that the different behaviour may be due to the fact that topologies are closely related to transcription, hence are expected to be cell-type specific, but the structural heterogeneity is not so easily captured by Hi-C like approaches. Second, while cluster formation (microphase separation) is the primary mode of transcriptional activation, several active genes are decompacted in our simulations, in line with experimental results showing increased enhancer-promoter distancing upon gene activation^16^. This conflict could reflect even decompacted genes having microphase separation in the vicinity of the promoter. However, the special case of extreme decompaction of highly active genes^34,36^ is not well modelled using the current framework, which would need to be modified to include, for instance, different levels of acetylation-related decompaction.

Epigenetic patterns differ widely between individual genes to drive changes in structure (Fig. 2), but epigenetic marks also vary within a single gene either between alleles or between individuals^37,38^. From the results discussed here, we would expect this to impact on both transcription levels but also transcription noise (heterogeneity), and consequently provide variation within the population which can exert evolutionary pressure. Additionally, alterations in epigenetic programmes that occur through genetic mutations could impact transcription in cells and tissues^39,40^, thereby providing a molecular basis for disease mechanisms.

## Acknowledgements

We would like to thank Ewan McDowall for developing the 3DGene database, Craig Nicol for designing the web page and the Edinburgh Compute and Data Facility (ECDF; http://www.ecdf.ed.ac.uk/). Thanks also to our group members and colleagues, in particular Javier Caceres, Martin Taylor and Jim Allan who provided advice during the project and comments on the manuscript. This work was funded by the European Research Council (ERC CoG 648050 THREEDCELLPHYSICS), UK Medical Research Council (MR/J00913X/1; MC_UU_00007/13) and the Wellcome Trust (223097/Z/21/Z).

## Author contributions

M.C., C.A.B., N.G., D.M. designed research; M.C., C.A.B., and C.B. performed simulations; C.N. and R-S.N. performed lab-based research; M.C., C.A.B., N.G., D.M. wrote the manuscript with input from all the authors.

## Data availability

The datasets generated during and/or analysed during the current study will be deposited on the Edinburgh University DataShare. To compare the predicted transcriptional activity to experiments, we used GRO-seq data GEO: GSM1480326. To relate simulated transcriptional noise to experimental data a PBMC3k dataset freely available from 10X Genomics was used.

## Code availability

The code used for the simulation is LAMMPS, which is publicly available at https://lammps.sandia.gov/. Custom codes written to analyse data are available from the corresponding author upon request, or they can be downloaded from https://git.ecdf.ed.ac.uk/dmarendu/3d-gene-structures (access can be requested from the corresponding authors).

---

**Extended Data Fig. 1:**
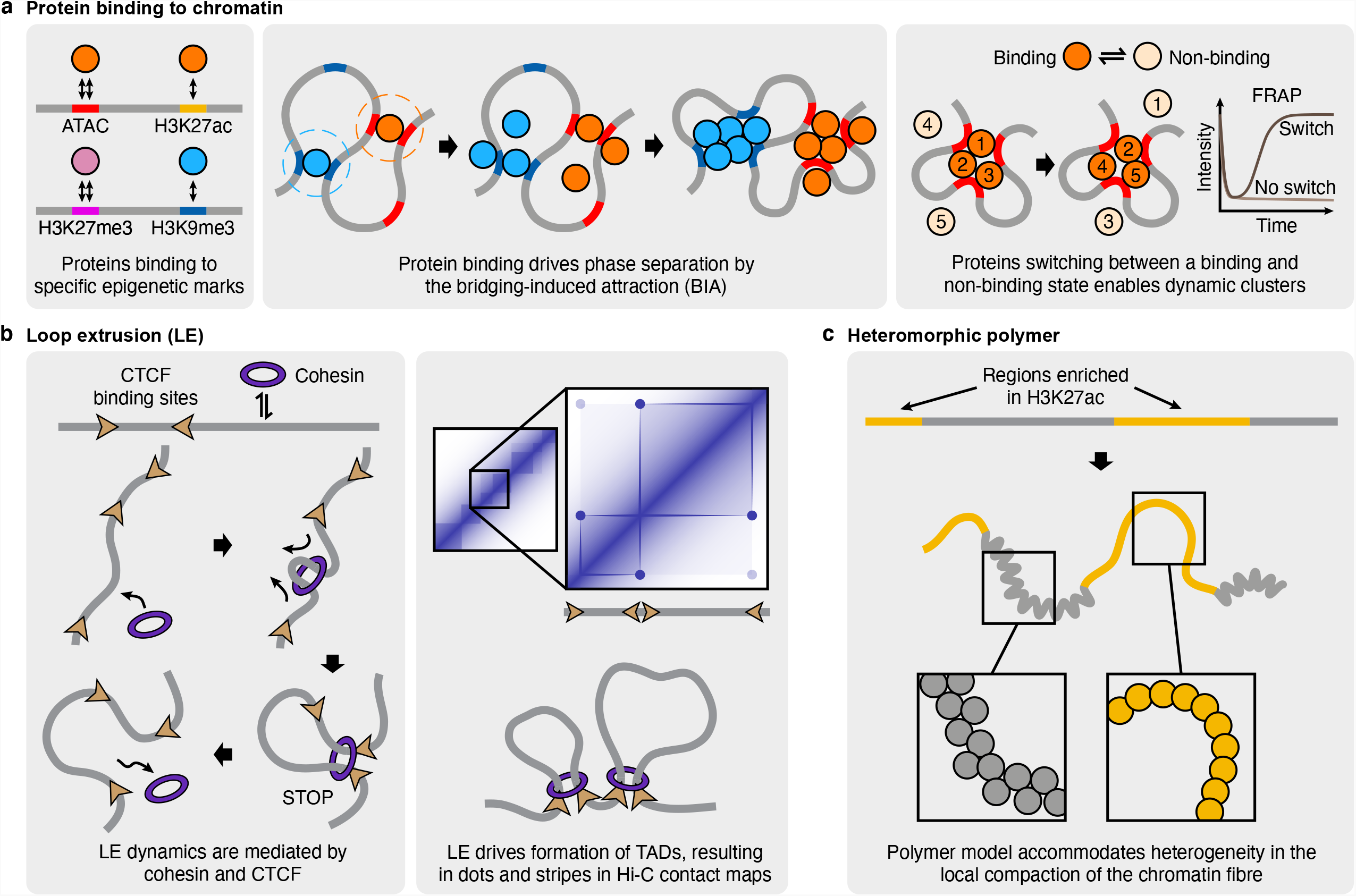
Key components of the HiP-HoP framework and their effect on chromatin structure and dynamics. **a**, The framework considers three species of proteins binding to chromatin loci enriched in specific epigenetic marks (left panel). With proteins being multivalent (and despite having no explicit attraction with one another), they drive the formation of protein clusters and chromatin domains via the bridging-induced attraction (middle panel): essentially, proteins bridge between chromatin loci sharing the same mark, increasing the local density of the mark, which in turn leads to more proteins attracted to the region, facilitating more bridging and a further increase in the density. This chain of events creates a positive feedback loop which ultimately allows proteins to cluster and phase separate, and concomitantly chromatin domains enriched in specific marks to develop. By allowing proteins to switch between a binding and non-binding state (right panel), the internal constituents of protein condensates become dynamic, recapitulating results from fluorescence recovery after photobleaching (FRAP) experiments. **b**, The model incorporates factors such as cohesin (or other SMC complexes) which move divergently along chromatin to extrude loops, and stop at CTCF boundary elements oriented opposite to the direction of travel (left panel). This mechanism favors the formation of convergent CTCF loops and produces chromatin domains with structural features consistent with those seen in Hi-C contact maps (right panel; e.g., dots for loop anchors at CTCF binding sites). **c**, Variation in local chromatin structure is modelled by adding extra springs linking between next-to-nearest beads (see boxes) in regions depleted of H3K27ac. This enables the model to reproduce the local structural decompaction or disruption in actively transcribed chromatin regions as observed in experiments^5^.

**Extended Data Fig. 2:**
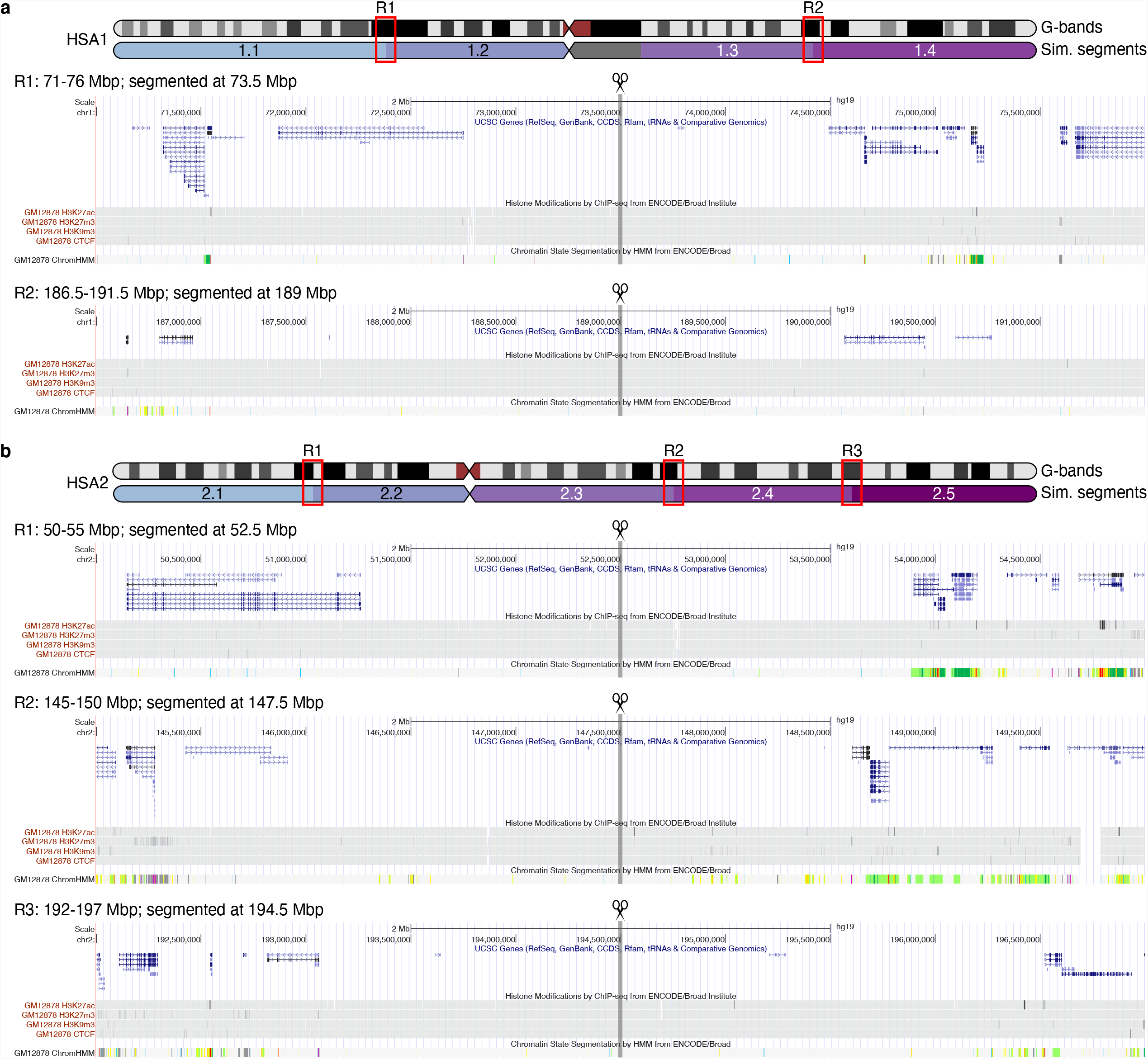
Long chromosomes are divided into shorter segments when performing HiP-HoP simulations. **a**, HSA1 was simulated as four separate segments. Ideograms at the top indicate the positions of segmentation (the region coloured in grey is unmappable within the hg19 reference genome and is therefore not simulated). Snapshots from the UCSC genome browser (bottom) show the genes, various ChIP-seq signals (H3K27ac, H3K27me3, H3K9me3 and CTCF) and the 15 chromatin-state hidden Markov model (HMM) track within a 5 Mbp region around two segmentation points, R1 and R2. These tracks demonstrate that there are no notable genomic and epigenomic features near the segmentation points. **b**, Similar to **a** but reports the segmentation points in HSA2 and the nearby chromatin context, with no notable features observed.

**Extended Data Fig. 3:**
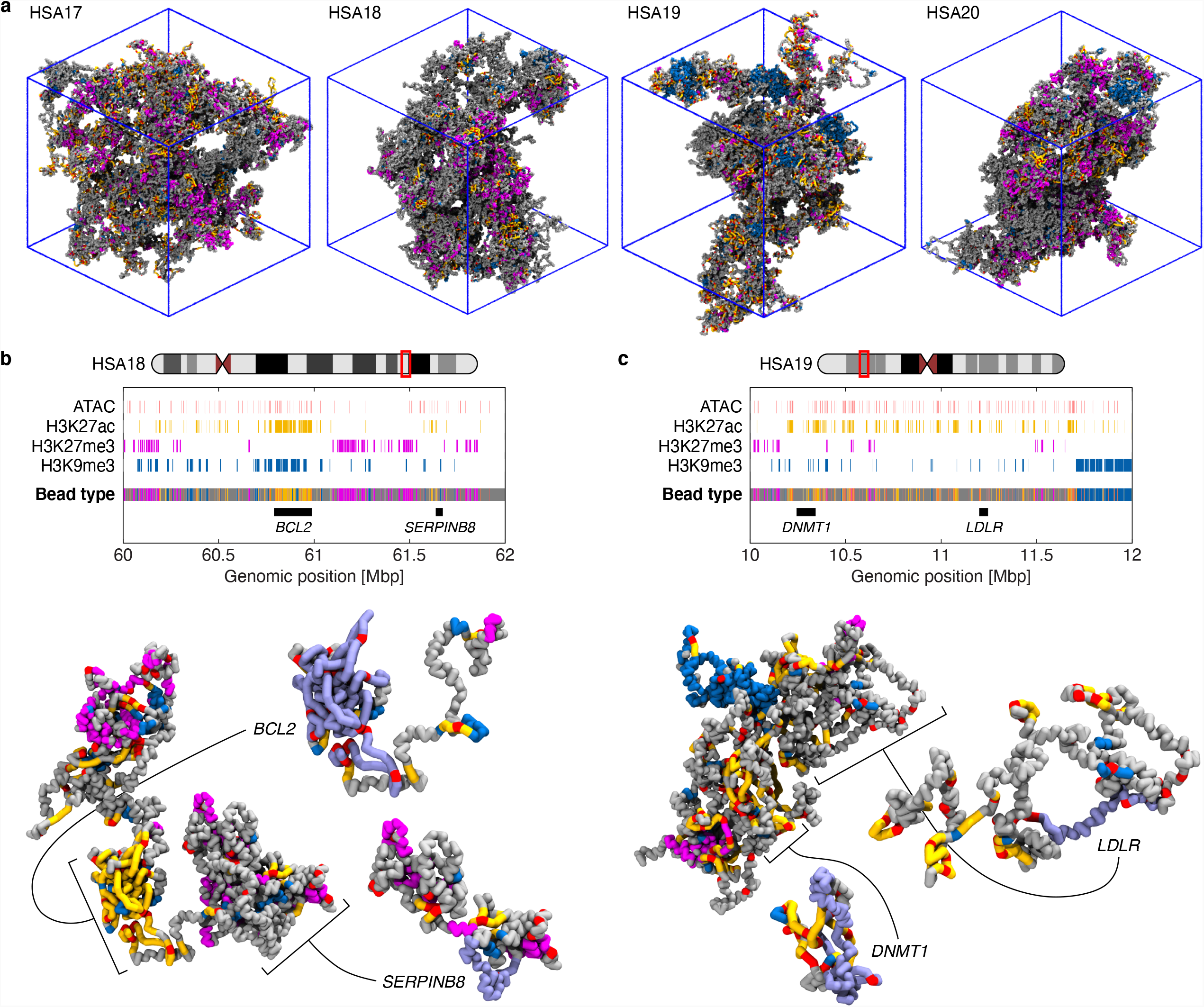
Simulated structures from HiP-HoP framework provide a detailed view of the 3D conformation of individual genes and their neighbourhood. **a**, Representative snapshots of the predicted structures of HSA17, 18, 19 and 20, all of which are simulated individually as a single, continuous polymer without segmentation. **b**, An enlarged view of the region 60-62 Mbp in HSA18. Epigenetic tracks used as model inputs and the resulting bead type along the chromatin fibre are shown at the top, and an example structure of this region is shown at the bottom, with a further expanded view of two gene loci, *BCL2* and *SERPINB8*. **c**, Similar to **b** but for the region 10-12 Mbp in HSA19, with the conformations of the gene loci *DNMT1* and *LDLR* shown.

**Extended Data Fig. 4:**
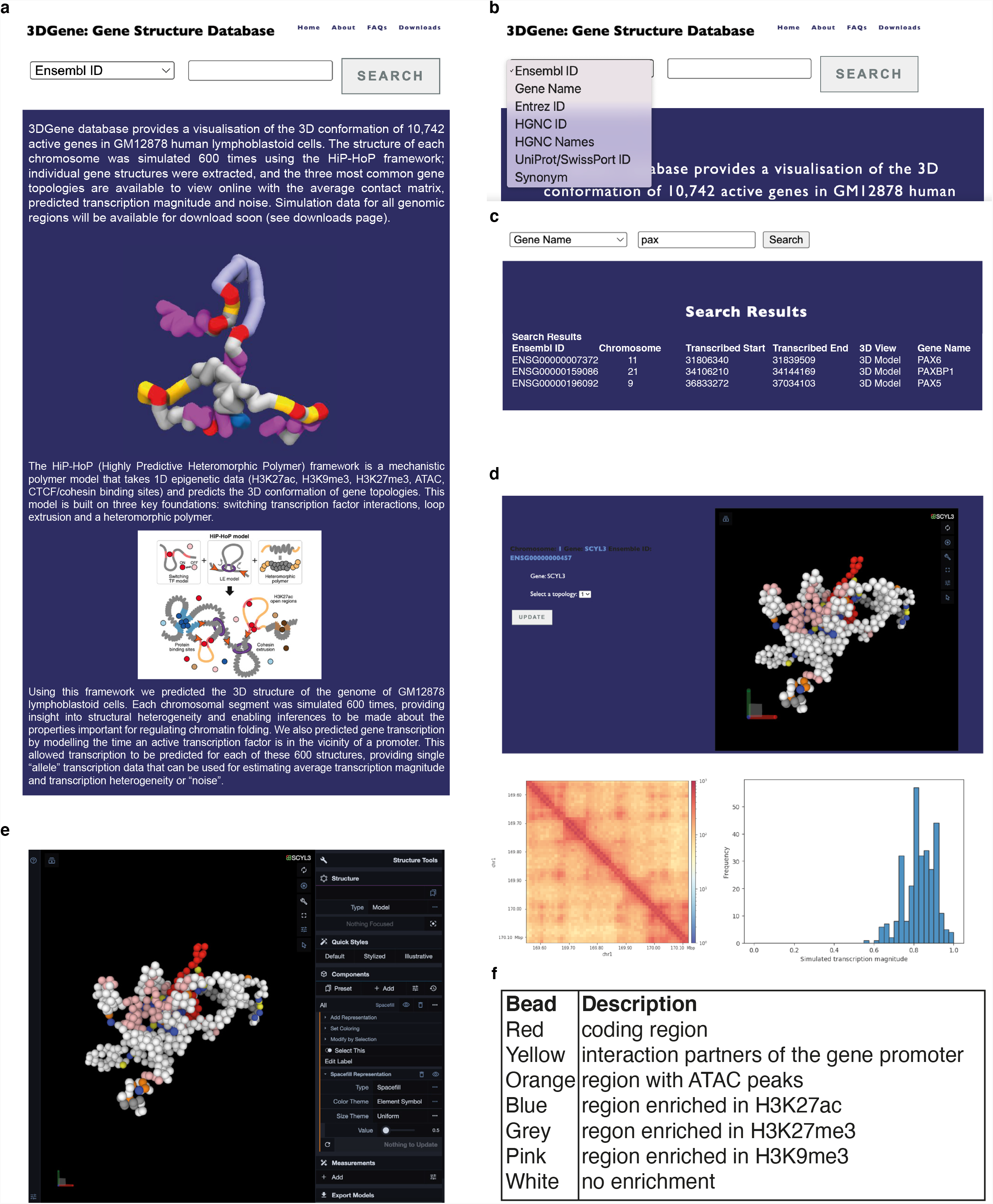
3DGene website provides a simple interface for browsing and downloading simulated gene structures from the 3DGene database. **a**, Interface to the ‘3DGene: gene structure database’ at https://3dgene.igc.ed.ac.uk. The database enables visualisation of 10,742 active genes in GM12878 human lymphoblastoid cells. **b**, Genes can be searched on common identifiers including Ensembl ID and Gene Name. **c**, After searching for a gene, using either a full or partial identifier, hits are returned with chromosome and gene co-ordinates (hg19). Selecting the 3D view reveals an interactive results page. **d**, The results page has four main elements. Gene details, interactive structure window, contact matrix and gene expression data. From the gene details panel one of the three most common topologies can be selected. The 3D structure of the topology is returned in the viewing panel. Each bead corresponds to 1 kbp of DNA and bead colours are representative of differently marked or ‘flavours’ of the chromatin fibre (see panel **f**). To the right of the viewing window there are controls. The square brackets maximise the window whilst the spanner reveals further visualisation settings (see panel **e**). The 3D structure can be rotated and zoomed in and out using the mouse; the aperture icon is used for capturing a screen grab. The contact matrix is a composite of all the interactions within the gene and is analogous to average Hi-C data from 300 cells (two alleles per cell). The transcription magnitude panel shows a histogram assembled from the simulated gene expression data for each different gene structure, akin to a single-cell sequencing experiment. The median is representative of average gene expression, whilst the width of the distribution is indicative of gene transcription heterogeneity or noise.

**Extended Data Fig. 5:**
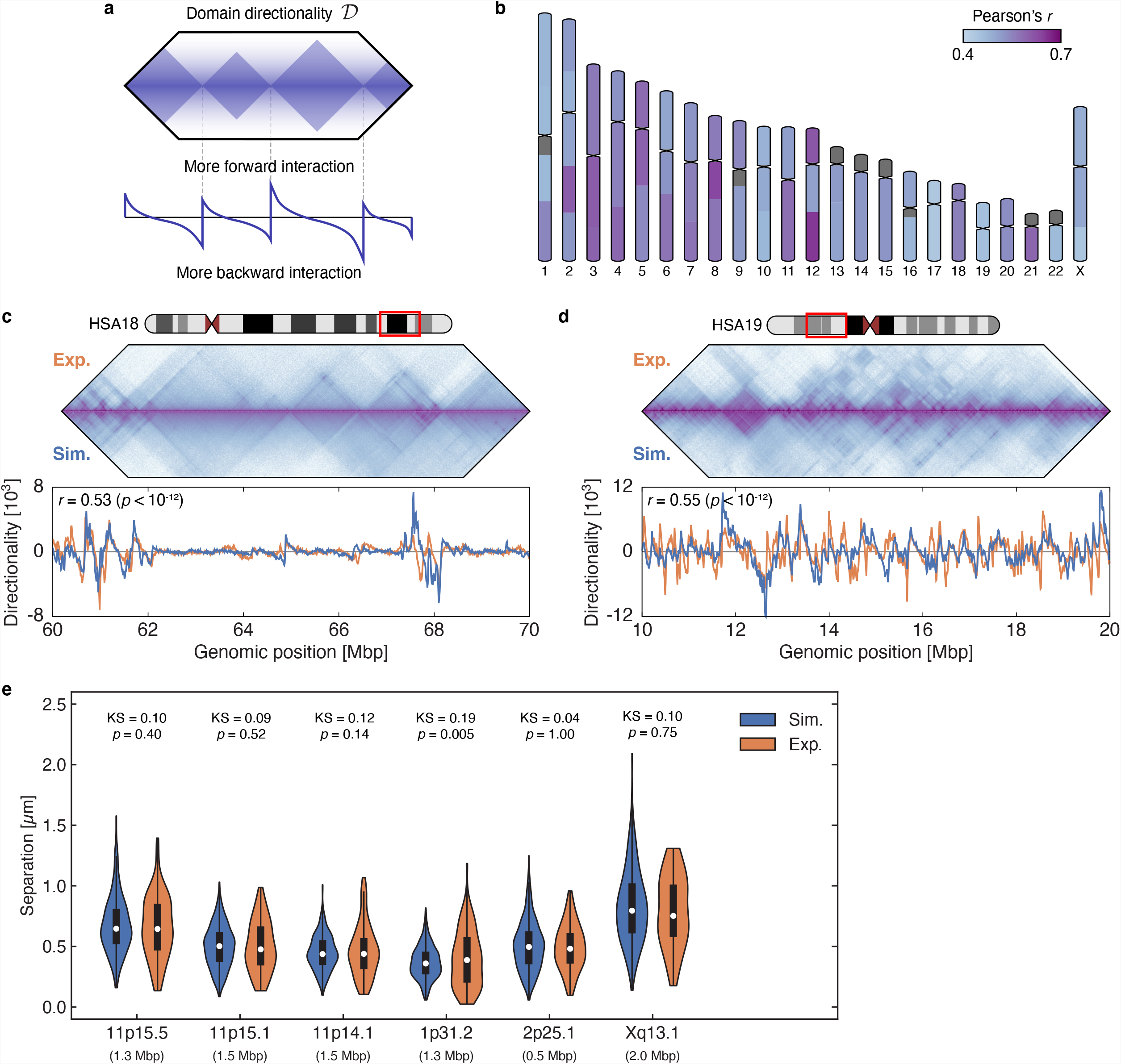
Validating the genome-wide HiP-HoP simulations with Hi-C and FISH data. **a**, Schematics explaining the directionality index *D* which quantitatively captures domains seen in contact maps. The *D* score at a chromatin bin is the total contact frequencies of the bin with other bins to its left, up to a threshold, subtracted by the total contact frequencies with those to its right (see Methods). **b**, Ideograms showing the Pearson correlation between the directionality profiles from simulated and Hi-C contact maps for each chromosome segment (grey regions are not simulated as they are unmappable regions within the reference genome). **c**, An example comparison between simulated and Hi-C contact maps for the region 60-70 Mbp in HSA18 (top) and between their directionality profiles (bottom). **d**, Similar to **c** but for the region 10-20 Mbp in HSA19. **e**, Comparing the simulated and experimental FISH distance distributions for six pairs of probes. As for Fig. 1i, we tested whether the difference between the distributions is statistically significant using the two-sample Kolmogorov-Smirnov (KS) test. Of all, the simulated and experimental distributions only show significant deviation for the probe pair within 1p31.2, which is a gene-poor region and is therefore less relevant to our analysis on actively transcribed chromatin. Experimental data are taken from Ref. ^43^.

**Extended Data Fig. 6:**
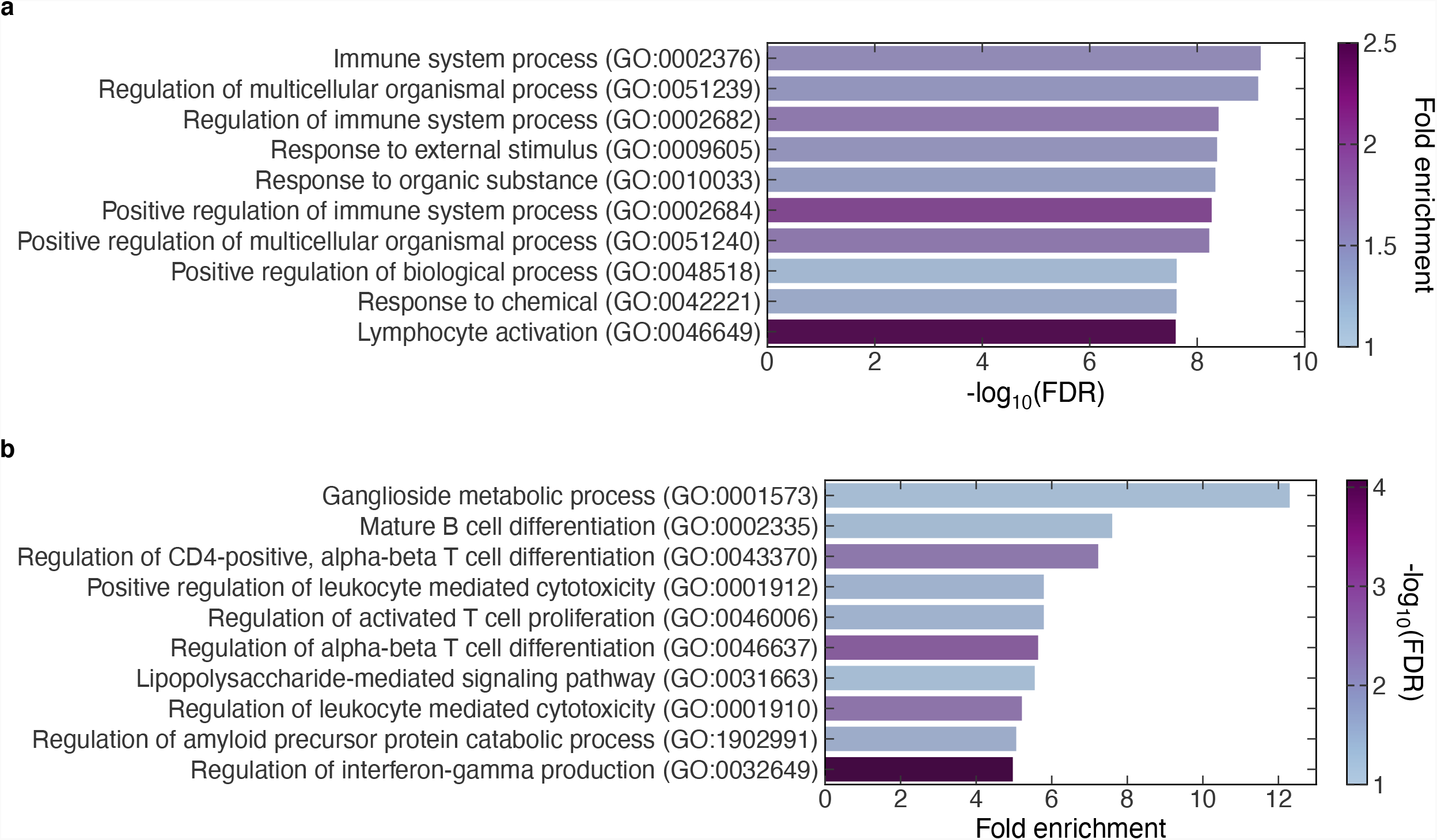
Gene ontology (GO) overrepresentation analyses reveal functional significance of genes with influential nodes. **a**, A list of the top ten GO biological function terms, ranked by -log_10_ of the false discovery rate (FDR), from performing an overrepresentation test in GO terms comparing genes with more than one influential node to those with only a single node. Test was completed using the web tool PANTHER (released date: 2022/02/02) and the GO database released on 2021/11/16. Most of these GO terms are related to immune response: since we simulated lymphoblastoid cells, this result indicates that genes with a higher number of influential nodes are typically more tissue specific. **b**, The top ten GO terms ranked by fold enrichment for the same overrepresentation analysis as done in **a**.

## Methods

### Polymer physics modelling

The HiP-HoP framework consists in coarse-grained molecular dynamics simulations, in which collections of molecules are represented by beads, which interact with phenomenological force fields and move according to Newton’s laws^5^. More specifically, chromatin fibres and chromosomes were modelled as bead-and-spring polymers, while complexes of transcription factors (TFs) or other chromatin-binding proteins were represented by additional individual beads. We also included loop extruding factors, represented by additional springs between non-adjacent chromatin chain beads, and chromatin heteromorphism. We used the multi-purpose molecular dynamics package LAMMPS^44^ (Large-scale Atomic/Molecular Massively Parallel Simulator). In this section we detail the potentials underlying the force fields used in the simulations.

#### The chromatin fibre

A chromatin fibre corresponding to a whole chromosome, or chromosome fragment, was discretised as a set of monomers, each of size corresponding to 1 kbp, or to *σ*∼20-30 nm, which can be determined by fitting to microscopy experiments (see below). Any two monomers (*i* and *j*) in the chromatin fibre interact purely repulsively, via a Weeks-Chandler-Anderson (WCA) potential, given by

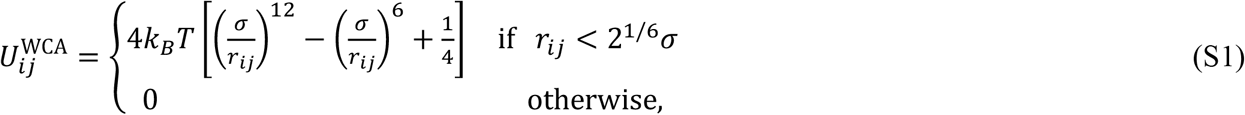

where *r*_*ij*_ is the separation of beads *i* and *j*. There is also a finite extensible non-linear elastic (FENE) spring acting between consecutive beads in the chain to enforce chain connectivity. This is given by

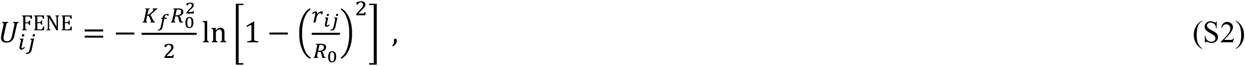

where *i* and *j* are neighbouring beads, *R*_0_ = 1.6 *σ* is the maximum separation between the beads, and *K*_*f*_ = 30 *k*_*B*_*T*/ *σ*^2^ is the spring constant. Additionally, a triplet of neighbouring beads in the chromatin fibre interact via a Kratky-Porod term to model the stiffness of the chromatin fibre, which explicitly reads as follows,

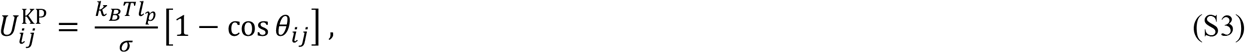

where *i* and *j* are neighbouring beads, while *θ*_*ij*_ denotes the angle between the tangent vector connecting beads *i* and *j = i+1* and that connecting beads *j* and *j+1*. The quantity *l*_*p*_ is related to the persistent length of the chain: this parameter is set to 4 *σ* in our simulation, which corresponds to a relatively flexible chain. Note however that the effective persistence length of the polymer can differ from the parameter value, as it depends on the local chromatin environment and its heteromorphic compaction.

#### Multivalent chromatin-binding proteins

Multivalent TFs are modelled as spheres, again with size *σ* for simplicity. The interaction between a chromatin bead, *a*, and a multivalent TF, *b*, is modelled via a truncated and shifted Lennard-Jones potential, given by

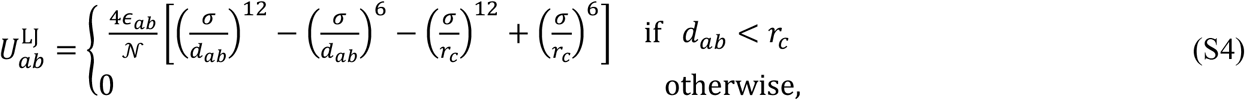

where *d*_*ab*_ denotes the distance between the centres of the chromatin bead and protein, *r*_*c*_ = 1.8 *σ* is a cut-off parameter, and *𝒩* is a normalisation constant which ensures the depth of the potential reaches −*ϵ*_*ab*_ at the minimum point.

Three species of multivalent TFs are considered: a generic active TF and two inactive TFs, one modelling polycomb-like proteins and another modelling heterochromatin protein 1 (HP1)-like, or other heterochromatic, proteins. Active TFs bind strongly (*ϵ*_*ab*_ = 7 *k*_*B*_*T*) to beads with high accessibility (corresponding to ATAC peaks; for more details on this and on chromatin bead colouring, see ***Simulation input data***) and weakly (*ϵ*_*ab*_ = 3 *k*_*B*_*T*) to those enriched in H3K27ac. This aims to capture promoter-enhancer interactions and the formation of transcriptional domains^26^. For inactive TFs, polycomb-like TFs bind to beads enriched in H3K27me3 (*ϵ*_*ab*_ = 7 *k*_*B*_*T*), representing interactions mediated by polycomb repressive complexes^45^ (PRCs). Heterochromatic TFs bind to beads with H3K9me3 (*ϵ*_*ab*_ = 3 *k*_*B*_*T*), modelling bridging facilitated by HP1^46^. Importantly, TFs only interact with one another via steric repulsion [described by the WCA potential, Eq. (S1)]. Nevertheless, thanks to their ability to bridge between multiple chromatin beads with similar marks, TFs of different species tend to microphase separate and form individual clusters via the bridging-induced attraction^6^ (Extended Data Fig. 1a). In addition to TF-chromatin binding, there is also a weak, direct chromatin-chromatin interaction (*ϵ*_*ab*_ = 0.4 *k*_*B*_*T*) between beads which do not possess the active H3K27ac mark. This extra ingredient facilitates the phase separation between euchromatin and heterochromatin.

TFs also switch back and forward between a binding and a non-binding state with a rate *k*_sw_. This feature mimics post-translational modifications on protein complexes and accounts for the dynamical turnover of constituents within nuclear protein clusters^47^ (Extended Data Fig. 1a). When TFs are non-binding, they interact with chromatin beads via steric repulsion (modelled by the WCA potential).

#### Loop extrusion

Loop extruding factors, representing for instance the structural maintenance of chromosome (SMC) complex cohesin, were incorporated in the HiP-HoP framework in a way similar to previous work^7^. Each loop extruder was described as a dimer whose two ends moved divergently along the fibre. For simplicity, this process was modelled implicitly as a harmonic spring with short range WCA repulsion between the chromatin beads *i* and *j* to which the two ends of an extruder are instantaneously bound,

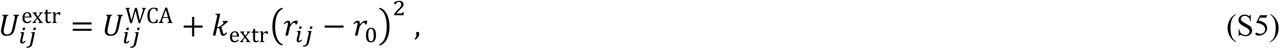

where *k*_extr_ = 40 (*k*_*B*_*T*)/ *σ*^2^ is the harmonic spring constant and *r*_0_ = 1.5 *σ* the harmonic bond length. In the simulations, an extruder could bind randomly to any chromatin bead, say the *i*-th one, and this was modelled by introducing a spring linking beads *i* and *i+3* (as a crumpled spring may already connect beads *i* and *i+2* to model the heteromorphic fibre, see below). Once bound, the two ends of an extruder were assumed to translocate at speed *ν*_extr_, and this was done by moving the spring to the next pair of beads. Both ends of an extruder moved along the fibre until colliding another extruder or reaching a CTCF bead (see below) with opposite orientation to the direction of travel. Note that the two ends move independently: if one end halts due to the aforementioned scenarios, the other can continue to extrude. Extruders detached from chromatin with rate *k*_off_, upon which the spring was removed. For simplicity, it was assumed that when an extruder unbinds from the fibre, another immediately re-attaches to it (i.e., the number of extruders on chromatin remains constant). The loop extrusion dynamics were performed using a python script which drives the LAMMPS library.

#### Heteromorphic chromatin modelling

A prominent feature of the HiP-HoP model is that it represents chromatin as a heteromorphic polymer, whose linear compaction varies along the contour. This feature is needed to account for a more open and disrupted conformation in acetylated regions, as observed in fluorescence in situ hybridisation (FISH) experiments^5^. In practice, the heteromorphic property was implemented by introducing extra harmonic springs which link next-to-nearest neighbour beads along the chain, *i* and *i+2*, in regions that are not annotated with the H3K27ac mark. Here, the spring constant was set to *k*_*h*_ = 200 *k*_*B*_*T*/ *σ*^2^ and the bond length to *r*_0_ = 1.1 *σ*. In this way, regions with the extra springs become crumpled and have higher linear compaction, whereas those with the mark remain open. This variability in local folding changes the stiffness and persistence length of the fibre^5^.

#### Equations of motion

The time evolution of each bead *i* in the simulation (whether TF or chromatin bead) is governed by the following Langevin equation:

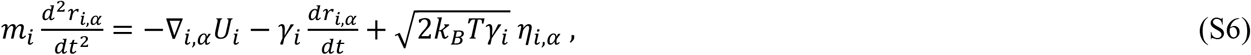

where *α* denotes Cartesian vector components, *r*_*i, α*_ is the *α*-th component of the position vector of bead *i, U*_*i*_ is the total potential experienced by it, *m*_*i*_ and *γ*_*i*_ are its mass and friction coefficients (these coefficients are the same for all beads in our simulations, denoted by *m* and *γ*), and *η*_*i, α*_ is the *α*-th component of a stochastic noise vector with the following mean and variance:

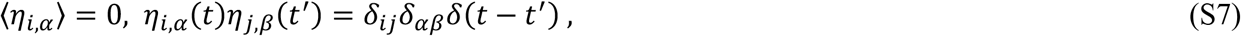

where the Latin and Greek indices run over particles and Cartesian components, respectively, and the first two *δ*’s denote Kronecker deltas, while the last one a Dirac delta.

### Simulation units and parameters

Our basic unit of length is the chromatin bead size *σ*, which sets the resolution and equals 1 kbp. Since the packaging of DNA into chromatin is not fully understood, we did not fix the physical value of the length unit in nm ahead of running the simulations. Instead, we performed the simulations and estimated the physical value of *σ* by comparing with experimental data, typically FISH (see below).

Regarding timescales, we note that the previously defined simulation units give rise a natural time unit, 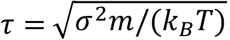. Another important timescale is the Brownian time *τ*_*B*_ = *σ*^2^/*D*, with *D* = *k*_*B*_*T*/*γ* the diffusion coefficient of the beads. The Brownian time gives an order-of-estimate measure of the time it takes for a bead to diffuse across its own diameter, and it is this timescale which we use to determine the mapping of simulation time to real time. In simulation units, for simplicity we work with all beads having the same mass *m* = 1 and set *γ* = 2, so that *τ*_*B*_ = 2*τ*. This means that the system is overdamped, as should be the case physically; however, beads have more inertia than in reality (this is necessary to keep overall simulation times manageable), but this will only affect very early times, whereas we are interested in chromatin configurations at steady state. To map between simulation and real time, we measured the average mean squared displacement for all polymer beads and found the value of *τ*_*B*_ which gives the best fit to experimental results from Ref. ^48^. By doing so, we got *τ*_*B*_ ∼ 5 ms. Numerical integration of Eq. (S6) was performed using a standard velocity-Verlet algorithm with time step *dt* = 0.01 *τ*_*B*_, implemented in the LAMMPS engine^44^.

Parameter values used in the simulations and not specified in the ***Polymer physics modellin****g* section above were chosen as follows. The number of TF beads was set to 10% of the number of chromatin beads, while the ratio of active, polycomb-like, and heterochromatic TFs was fixed at 1/4:1/8:5/8. Additionally, the switching rate was set to *k*_sw_ = 10^−3^*τ*^−1^. As for loop extrusion, the number of extruders bound to chromatin was fixed to give a density of 10 extruders/Mbp. Extrusion and unbinding rates were set to *ν*_extr_ = 4 × 10^−3^ Kbp/*τ* and *k*_off_ = 2.5 × 10^−5^*τ*^−1^, respectively. While these parameters were chosen to facilitate sampling and are less realistic individually, the ratio *λ* = *ν*_extr_/*k*_off_ = 160 kbp, also known as the extruder’s processivity, and the density of extruders (or the average spacing between them) are consistent with values used in the literature^7^.

The chromatin fibre was simulated within a periodic cube of length *L* such that the density of chromatin is ∼6.5 Mbp/μm^3^. This is based on the fact that there are around 6.5 Gbp of DNA in a human diploid cell, and that the diameter of a typical cell nucleus is ∼10 μm.

To facilitate computation, chromosomes were simulated individually rather than as a whole (Fig. 1c and Extended Data Fig. 2). While this approach neglects long-range and interchromosomal (or *trans*-) interactions, this is acceptable in our case as the main objective of the work is to examine the local chromatin structure and *cis*-interactions of regulatory elements, which typically span no longer than a few Mbps. In practice, shorter chromosomes (chromosomes 14, 15 and 17 to 22) were simulated as a single segment, whereas longer chromosomes (chromosomes 1 to 13, 16 and X) were each divided into smaller segments. Breakpoints were chosen to be sites where there is low enrichment in chromatin modifications within their neighbourhood (i.e., gene deserts; see Extended Data Fig. 2).

### Initialisation and simulation protocol

The simulation system was initialised as follows. As described in Refs. ^5,49^, the chromatin fibre was first generated as a mitotic-like helix conformation – i.e., a stack of rosettes (see Ref. ^49^ for details on the equations used to create these). The initial simulation box was a parallelepiped, with equal smaller dimensions in the *x* and *y* directions and a longer height that depended on the length of the initially cylindrical chromosome segment along *z*. To relax the fibre from the rosette configuration, beads along the chain were initially connected together using harmonic springs (with spring constant equal to 100 *k*_*B*_*T*/ *σ*^2^ and bond length equal to 1.1 *σ*), and all non-neighbour pairwise interactions were governed by the repulsive soft potential (whose strength gradually increased from 0 *to* 500 *k*_*B*_*T*). After an initial relaxation simulation of duration 500 *τ*, the springs were replaced by FENE bonds and the soft potential by the WCA potential. The simulation box was then compressed slowly along the *z* direction for 10^4^*τ* such that it became a cube with the desired volume. Next, the fibre was relaxed further within the cube with fixed boundaries for 5 × 10^3^*τ* and then with periodic boundaries for 2.5 × 10^4^*τ*. To allow the fibre to quickly lose memory of the rosette-like conformation, extruders (with a density of ∼7.5 extruders/Mbp) were loaded to the fibre to perform loop extrusion without any CTCF boundaries for 2 × 10^4^*τ*. Harmonic springs for modelling fibre heteromorphism were then added to the entire fibre, and the fibre was allowed to relax for 10^4^*τ*. Next, TF beads were incorporated and allowed to equilibrate with the fibre for 10^3^*τ*, with TF-chromatin interactions being purely repulsive (modelled by the WCA potential). Finally, regions enriched in H3K27ac had their crumple springs removed such that the fibre had different levels of local compaction.

10 independent runs were conducted using this procedure for each chromosome segment, and their final conformations were used as the starting conditions for 300 production runs (which were performed using different random seeds). From measuring structural properties such as the radius of gyration of the fibre, it was verified that production runs starting from the same relaxed conformation gave different structures. In the production run, the chromatin fibre was simulated for a period of 3 × 10^5^*τ*. Attractive chromatin-chromatin interactions were switched on at time 10^4^*τ*, and subsequently TF-chromatin interactions at 5 × 10^4^*τ*. The system was sampled (e.g., for contact map calculation) every 2 × 10^3^*τ* during the final 10^5^*τ* of the simulation period. The 600 structures shown in the main text were generated at times 2 × 10^3^*τ* and 3 × 10^5^*τ*.

### Simulation input data

Chromatin beads were assigned different states or colours according to their local epigenetic modifications and DNA accessibility. Three epigenetic modifications have been used here: H3K27ac, H3K27me3 and H3K9me3. These marks are typically associated with actively transcribed euchromatin, facultative heterochromatin and constitutive heterochromatin, respectively. Their ChIP-seq profiles (for GM12878 cells, aligned to the hg19 reference genome) are obtained from the ENCODE database (https://www.encodeproject.org). H3K27ac data were used to determine disrupted regions of the chromatin fibre, where no additional crumpling springs were applied; H3K27me3 and H3K9me3 tracks were used to set binding sites for polycomb-like and heterochromatic proteins. DNA accessibility was determined from an assay for transposase-accessible chromatin using sequencing (ATAC-seq) data set^50^. ATAC-seq peaks were used to identify high-affinity binding sites for multivalent active proteins. Beads were assigned particular epigenetic and/or ATAC marks if the respective chromatin regions have significant enrichment, or peaks, of these marks. Note that a bead could have more than one kind of mark, and its state or colour was determined based on the combination of marks it possesses.

To identify the CTCF binding sites which constrain loop extrusion, ChIP-seq data sets for CTCF and Rad21, a subunit of the cohesin complex, were obtained from ENCODE. Genomic loci which had peaks in both profiles while also containing the CTCF binding motif were marked as candidate binding sites. Beads covering these sites were then labelled as CTCF beads, and the direction in which they act on the extruders was based on the orientation of the underlying motif. Motivated by the cell-to-cell variability in CTCF binding, CTCF beads were activated stochastically in each simulation run according to a probability that was linearly proportional to the score of the corresponding CTCF peak, similarly to Ref. ^5^. When there were multiple peaks within the same bead, all possible outcomes were considered. For example, if a bead encompassed both a forward and backward-oriented CTCF binding site, the probabilities of the bead being a forward, backward, bidirectional or inactive CTCF boundary were calculated based on the score of individual peaks, and an outcome was selected based on these probabilities.

### Comparison with Hi-C, GRO-seq, FISH, and scRNA-seq experiments

Before HiP-HoP could be used genome-wide in a predictive way, it was important to validate it by showing that it gives results consistent with experimental data. To do so, HiP-HoP predictions were compared to four types of experiments: two measuring structure (Hi-C and FISH), and two measuring transcription (GRO-seq and single-cell RNA-seq). Below we describe each of these comparisons in turn.

#### Hi-C contact maps

To assess the ability of HiP-HoP simulations to predict structural data, contact maps were generated from simulations and compared to those from Hi-C experiments^15^. Simulated maps were constructed in a way inspired by the cross-linking step in Hi-C for sampling pairwise chromatin interactions^5^. As previously mentioned, the conformation of the chromatin fibre was sampled every 2 × 10^3^*τ* over a period of 10^5^*τ*. For each sampled structure, the following procedure is conducted. First, a pair of chromatin beads, say *i* and *j*, were selected randomly with their separation *r*_*ij*_ computed. Then, the pair was accepted and registered as a ‘read’ in the contact matrix with probability

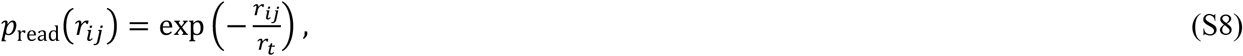

where *r*_*t*_ is a distance threshold related to the separation within which crosslinking is effective. By repeating this procedure many times and over different structures and simulation runs, reads can be piled up in a way similar to that in Hi-C. By fixing the total number of reads in the simulated map to be the same as that in a Hi-C map, one can directly compare the two without performing rescaling.

Experimental and simulated contact maps can be qualitatively compared by plotting them together, as in Fig. 1g of the main text. To quantify the similarity between maps, a simple way would be to compute the Pearson correlation of the maps directly; however, since the interaction probability between two loci is strongly dependent on their genomic distance, a direct correlation is usually not informative, as the resulting score predominately reflects this dependence. In particular, correlating two maps with very different interaction patterns can still achieve a reasonable score. Hence, a robust method should factor out this distance dependence while still having the power to discern differences between maps. A “distance-insensitive” strategy to evaluate the agreement between simulated and Hi-C maps is to compare the pattern of contacts at the scale of topologically associating domains (TADs) between them. One approach to detect TADs algorithmically from a contact map is to consider the fact that near their boundaries, interactions are highly favoured towards either the left or right of a chromatin segment^35^. This bias is captured by a directionality score *D*(*i*) (Fig. 1g and Extended Data Fig. 5a), which is defined for each chromatin bin (say bin *i*) in the map by subtracting the interactions in the region to the left of *i* by the interactions to the right of *i*. For both right and left interactions, contacts were summed up between a lower and an upper end threshold. The lower end threshold was fixed at 20 kbp to avoid artifacts close the diagonal within the Hi-C map (which were considered at 10 kbp resolution), whereas the upper end threshold was set at 500 kbp (varying between 100 kbp and 2 Mbp showed no significant differences). Note that because this score compares the aggregated interactions within regions either side of a bin up to the same genomic distance away, the distance dependence is effectively cancelled out. The similarity between the simulated map and the Hi-C map can then be determined based on the Pearson correlation of *D*(*i*) in both maps (Fig. 1g and Extended Data Figs. 5b-d).

Parameters listed in the ***Simulation units and parameters*** section were chosen so as to optimise the comparison between simulated and Hi-C contact maps for the genomic region HSA19: 6.5–16.5 Mbp (a 10000-bead segment). There are two main reasons for selecting this region. First, since this work focuses on the spatial interactions between regulatory elements, HSA19 is an ideal candidate as it has a high gene density and thus provides more data per base pair than other chromosomes. Second, the specified region still contains both transcriptionally active and inactive domains, so it is possible to determine the effects of different TF species.

#### Comparison with FISH experiments

Another way to compare predicted and observed 3D chromatin structure is to simulate FISH experiments. To do so, three pairs of FISH probes were considered from experiments using SATO3 lymphoblastoid cell ^41^ (^41^Figs. 1h,i), together with six pairs of probes from experiments using RPE1 cells^43^ (Extended Data Fig. 5e). For each pair, to generate probe separation measurements from our simulation data, we identified chromatin beads associated with each probe and took its position to be the centre of mass of these beads. By measuring the separation of the centres of mass of beads mapping to the pair of probes in each of the 600 simulated structures, a distribution of separations, in units of *σ*, can be constructed.

This simulated distribution can then be compared to the corresponding experimental one. To perform this comparison quantitatively, we mapped simulation length units to physical ones. For each pair, we used the two-sample Kolmogorov-Smirnov statistics to determine the distance between the simulated distribution and the corresponding experimental one, and found the value of *σ* in nm which minimises this distance. We then computed the *p* value corresponding to the probability that the experimental and simulated distributions were the same. This procedure identified that experimental and simulated distributions were not different in a statistically significant way (*p* > 0.1) in all but one case (see Extended Data Fig. 5e). This case corresponded to a gene-poor region (1p31.2), which may be less accurately represented by our simulations. Additionally, experimental measurements in this case were based on RPE1 cells, which may have a different local environment in that region with respect to the GM12878 lymphoblastoid cells that we simulated.

#### GRO-seq and patterns of transcriptional activities

To predict transcriptional activity of each ATAC bead in our model, we hypothesised that the frequency at which an ATAC bead is bound by (i.e., within a distance of 3.5 *σ* from) one or more active TFs was monotonically related to the transcriptional output of the corresponding chromatin segment, with a higher frequency associated with higher output (Fig. 3a). In this way, 300 values of transcriptional activity (one per simulation) for each ATAC bead were generated and used to calculate distributions (one per ATAC bead). From these distributions, averages, with the meaning of average transcriptional activity of ATAC beads, could be compared genome-wide with experimental data. In particular, data from global run-on sequencing (GRO-seq) were used ^51^, as they provided a genome-wide, per-base measure on this activity.

#### Single-cell RNA-seq and transcriptional noise

HiP-HoP predictions for transcriptional noise are more challenging to experimentally validate genome-wide due to scarcity of data on non-genic nascent transcription in single cells. As a proof of principle, predicted levels of transcriptional noise were compared to single-cell RNA-seq (scRNA-seq) data on transcription in peripheral blood mononuclear cells (PBMC), which should be related, though not identical, to the GM12878 lymphoblastoid line. The PBMC scRNA-seq data set used is freely available from 10X Genomics and is the same as that analysed in the online tutorial at https://satijalab.org/seurat/articles/pbmc3k_tutorial.html. The data set provides transcriptome profiles from 2,700 single cells which were sequenced on the Illumina NextSeq 500.

To compare simulated and experimental transcriptional noise at gene promoters, data pre-processing and quality control were performed as detailed in the tutorial above, with the count matrix normalised such that the sum of counts for each cell is the same. Experimental data for genes with an average transcriptional activity larger than 10^−6^ were further filtered to eliminate cases with very sparse reads. For the remaining dataset, the coefficient of variation (CV; standard deviation over average) of transcriptional activity was computed and compared to the corresponding simulated value for each gene, resulting in a statistically significant agreement as shown in Fig. 4b in the main text.

### Analysis of 3D simulated chromatin structure

#### Identification of topologies

Within the main text, partners of a promoter were defined as ATAC beads which contact that promoter in over 10% of the simulated structures, and a gene locus defined as the union set of a promoter and all its partners. Given these definitions, one can determine the interaction topologies within a locus, or the different ways that the promoter of interest networks with its partners (Fig. 2e). Fig. 2f in the main text displays a plot of the number of topologies against the number of partners for all gene loci, showing that the former quantity increases rapidly with the latter, until saturating due to the limited number of conformations. This increase, to a first approximation, can be understood from a simple combinatorial calculation. If one neglects the polymeric nature of chromatin, finding the number of unique topologies for *q* partners is equivalent to counting the number of ways to create a subset within a set of *q* (distinguishable) elements. As each element can either be in the subset or not, this number is simply 2^*q*^. Clearly, this constitutes the maximum possible number of topologies, as physical constraints of the fibre would render some not achievable. Additionally, it is not clear *a priori* whether all the remaining permissible topologies would occur in reality, as other factors, such as the local chromatin context and linear spacing between ATAC sites, can also influence the interaction networks. Nevertheless, Fig. 2f shows that even with a modest number of partners, many topologies are detected for individual ATAC loci, suggesting that these loci display large heterogeneity in their 3D folding across the ensemble of simulated structures. This result is consistent with recent single-cell studies revealing the extensive variation in chromatin organisation within a population of cells^52^.

#### Shannon entropy and structural diversity

The *SMARCA5* and *GINS4* loci discussed as examples in Figs. 2h-i demonstrate that the number of conformations associated with individual topologies can vary substantially. In *SMARCA5*, many conformations are mapped to a single topology (Fig. 2h), whereas in *GINS4*, they are spread more evenly across a larger number of topologies (Fig. 2i). How the population of structures is distributed among the observed topologies can be quantified by a structural diversity score *H*, which is inspired by the Shannon diversity index or Shannon entropy. This quantity is defined as follows,

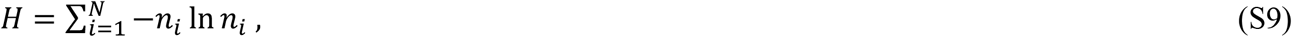

where *n*_*i*_ is the fraction of structures in topology *i*, and *N* denotes the total number of topologies. Note that *H* is expected to scale linearly with the number of partners *q*. This can be seen for instance from maximising *H*, which is achieved when *n*_*i*_ = 1/*N*. In this case, *H* = ln(*N*), and substituting the upper bound *N* = 2^*q*^ gives *H* = *q* ln(2). For this reason, the most informative genome-wide correlations described in the text were found with normalised structural diversity (equal to *H*/*q*).

